# APOBEC Mutagenesis is Concordant Between Tumor and Viral Genomes in HPV Positive Head and Neck Squamous Cell Carcinoma

**DOI:** 10.1101/2021.02.27.433168

**Authors:** Daniel L Faden, Krystle A. Lang Kuhs, Maoxuan Lin, Adam Langenbucher, Maisa Pinheiro, Meredith Yeager, Michael Cullen, Joseph F. Boland, Mia Steinberg, Sara Bass, James S. Lewis, Michael S Lawrence, Robert L Ferris, Lisa Mirabello

## Abstract

APOBEC (apolipoprotein B mRNA-editing enzyme, catalytic polypeptide-like) is a major mutagenic source in human papillomavirus positive oropharyngeal squamous cell carcinoma (HPV+ OPSCC). Why APOBEC mutations predominate in HPV+OPSCC remains an area of active investigation. Prevailing theories focus on APOBECs role as a viral restriction agent. APOBEC-induced mutations have been identified in both human cancers and HPV genomes, but whether they are directly linked in HPV+OPSCCs remains unknown. We performed sequencing of host somatic exomes, transcriptomes and HPV16 genomes from 79 HPV+ OPSCC samples, quantifying APOBEC mutational burden and activity in both the host and virus. APOBEC was the dominant mutational signature in somatic exomes. APOBEC vulnerable PIK3CA hotspot mutations were exclusively present in APOBEC enriched samples. In viral genomes, there was a mean (range) of 5 (0-29) mutations per genome. Mean (range) of APOBEC mutations in the viral genomes was 1 (0-5). Viral APOBEC mutations, compared to non-APOBEC mutations, were more likely to be low-variant allele frequency mutations, suggesting that APOBEC mutagenesis is actively occurring in viral genomes during infection. Paired host and viral analyses revealed that APOBEC-enriched tumor samples had higher viral APOBEC mutation rates (*p*=0.028), and APOBEC-associated RNA editing (p=0.008) suggesting that APOBEC mutagenesis in host and viral genomes are directly linked. Using paired sequencing of host somatic exomes, transcriptomes, and viral genomes from HPV+OPSCC samples, here, we show concordance between tumor and viral APOBEC mutagenesis, suggesting that APOBEC-mediated viral restriction results in off-target host-genome mutations. These data provide a missing link connecting APOBEC mutagenesis in host and virus and support a common mechanism driving APOBEC dysregulation.

## Introduction

The apolipoprotein-B mRNA-editing catalytic polypeptide-like (APOBEC) 3 family of cytidine deaminases is a major mutagenic source in human papillomavirus (HPV)-mediated cancers, including cervical and oropharyngeal squamous cell carcinoma (HPV+OPSCC). APOBEC-mediated mutations constitute a high proportion of the mutations in HPV+OPSCCS and can result in driver mutations such as activating mutations in PIK3CA^1,2^. The underlying mechanisms driving APOBEC dysregulation and mutagenesis in HPV+OPSCC remains an area of active investigation. One hypothesis focuses on APOBECs role as a viral restriction agent, with resultant “collateral damage” host genome mutations following APOBEC activation by viral infecton^3,4^. This hypothesis is supported by evidence that APOBEC: 1) acts to inhibit HPV both directly (with cytidine deamination of viral DNA leading to degradation) and indirectly (for example, through decreasing HPV virion infectivity^5^), 2) is upregulated both directly by HPV and by IFN as part of the innate immune response to viral infection^5-8^, 3) viral editing is identifiable and reproducible *in vitro* through APOBEC induction and in vivo in cervical pre-cancers and cancers^9-12^ and 4) mutations are present in nearly all human cancers but are particularly prominent in HPV+OPSCC, where they are associated with multiple measures of immune upregulation^13-22^.

Recently our group published a large case-control study of 5,328 cervical HPV genomes utilizing a HPV whole genome sequencing (WGS) approach, characterizing and annotating mutations across the HPV16 genome^23^. This study identified APOBEC-associated mutational signatures in the HPV16 genome and determined that APOBEC-induced viral mutations contributed to viral evolution and were significantly associated with reduced carcinogenicity of HPV16. While evidence supports the presence of APOBEC-induced mutations in both the genomes of HPV-mediated cancers and in HPV itself, whether APOBEC mutational activity in host and virus are directly linked remains unknown. Here, using 79 HPV+OPSCC samples with paired sequencing of host somatic exomes, transcriptomes and viral whole genomes, we apply computational approaches to characterize APOBEC activity in both host and viral genomes.

## Methods

### Samples

79 HPV+OPSCC tissue and paired blood samples were identified through existing databases at Vanderbilt University Medical Center and the University of Pittsburgh. HPV status was determined by p16 staining and subsequently by WGS for detection of viral genomes. After pathology review, DNA and RNA were extracted from FFPE tissue blocks using Zymo Quick DNA and RNA FFPE kits. Of these, 18 samples had been previously contributed to The Cancer Genome Atlas (TCGA) from the same institutions and had pre-existing whole exome sequencing (WES) and RNA-Seq. For these cases, DNA was extracted from FFPE blocks for viral WGS alone.

### Sequencing

Somatic DNA underwent WES using the KAPA HyperPrep Library Preparation kit followed by hybrid capture with the Illumina Rapid Capture Exome enrichment kit. All sample pairs were validated with Fluidigm fingerprint data to confirm sample identity and fidelity. Average (range) coverage was 218x (70-402x). Somatic RNA underwent RNA-Seq using the Illumina TruSeq RNA Exome kit. Average (range) number of reads per samples was 151M (17.5M-333M). HPV16 DNA underwent WGS as previously described by custom AmpliSeq™ panels followed by sequencing on an Ion Torrent platform with average (range) coverage of 23,000x (1,500x-56,000x)^24^. 74 WES-seq, 68 RNA-seq, and 72 viral WGS samples passed quality control. Of these samples, 63 had data available from all three platforms.

### Informatics

#### Somatic datasets

Somatic SNVs were called using MuTect v1. Mutations were screened against Panel of Normal (PoN) databases to remove common SNPs and recurrent sequencing artifacts. SNP-counting and copy number segmentation were done using FACETS. SNP counts were generated with minimum mapping quality of 15, minimum base quality of 20, pseudo-spacing of 100 and minimum read count of 25. Copy number data was segmented using window size of 1000, Variant Allele Frequency (VAF) threshold of 0.3 and cval of 300. Allelic amplifications/deletions were defined as regions of any size with rounded integer total copy number states above or below 2, respectively. Somatic mutation lists were combined with mutations lists from the entire TCGA cohort, and mutational signatures were deconvolved using NMF (K=8), as described previously.^19,25^ IFN-γ score was defined as the mean expression of six genes (IFNG, IDO1, CXCL9, CXCL10, HLA-DRA & STAT1) as described previously^26^. Analysis of PI3K/AKT/mTOR pathway was conducted using a six major effector gene signature (PI3K, AKT1, AKT2, AKT3, PTEN & MTOR) as described previously^27^, and further, hotspot loci were examined in IGV. Significantly mutated genes were identified using MutSig2CV, and significantly copy-number-altered genes were identified using GISTIC 2.0. MutSig and GISTIC p values were combined using Fisher’s method to calculate a unified p value per gene. Pathway enrichment scores were then calculated by combining unified p values for genes in a pathway, also using Fisher’s method.

#### Viral datasets

All HPV16 WGS data were aligned to a consensus HPV16 sequence NC_001526.4 from PaVE^28^. Position-wise base counts were obtained, and a major haplotype was created by taking the most prevalent nucleotide at each position. These haplotypes were fed into multi-alignment tool MUSCLE, along with multiple reference builds for HPV16 (AF402678, AF472509, AF534061, AF536179, AF536180, AY686579, HQ644236, HQ644257, HQ644298, K02718), and reference sequences for HPV35, HPV33, & HPV18. The resulting alignment was fed into PhyML (run using TOPALi v2.5), assuming a transversion model and allowing for invariable sites and specifying a gamma distribution. The resulting phylogenetic tree allowed for sublineage assignment of each sample by nearest distance to a reference sequence. All samples were realigned with respect to their relative reference. Putative mutation sites were nominated by having a minimum total coverage of 100, minimum alternate allele fraction of 0.02, and minimum alternate reads of 2. Putative mutations were anonymized, and manually reviewed in IGV alongside other samples aligned to the same reference to remove low-evidence mutations and recurrent sequencing artifacts.

Viral APOBEC-induced mutations were identified as C->T and C->G mutations at T<u>C</u>W motifs (W is A or T), as described previously^23^. To count the number of APOBEC targetable sites, we first counted TCW motifs across the HPV16 genome for each reference build and then the total number across all samples’ HPV16 genomes on their respective reference builds. Since there are three possible changes at each nucleotide position, APOBEC-targetable sites were counted as one-third of the total number of T<u>C</u>W motifs. Mutation rates were calculated as the total number of APOBEC mutations, divided by the total number of APOBEC-targetable sites at the gene level and genome level, stratified by mutation type (synonymous (S) and non-synonymous (NS)) and variant allele fraction (VAF) (low-VAF (VAF<= 0.5) and high-VAF (VAF > 0.5)).

## Results

### APOBEC mutations in somatic cancer genomes

79 HPV+OPSCC primary tumor samples were included in the analysis. The average patient age was 55. 82% of the cohort were male and 60% had a history of tobacco exposure of >1 pack year. Somatic exomes were pooled with TCGA data and mutational signatures extracted. As expected, HPV+OPSCC samples predominately clustered with other tumors dominated by the APOBEC mutational signature, with variable APOBEC mutational burden between samples (Figure 1, Supplemental Figure 1A). PIK3CA hotspot mutations, which have previously been described to be caused by APOBEC activity in HPV+OPSCC, were exclusively associated with APOBEC enriched samples (Figure 1B). The PI3K/AKT/mTOR pathway was disproportionately mutated across the cohort (p < 3.22×10^−8^) and amplifications in chromosomes 3q, which contains PIK3CA, were present in 41 samples. Both of these alterations have previously been described to occur frequently in HPV+OPSCC^17^. APOBEC mutations, A3A expression and APOBEC RNA editing, as measured by hotspot *DDOST*^*558C>U*^ mutations, were weakly correlated, consistent with prior reports, likely due to the episodic nature of APOBEC activity (Figure 1C).

**Figure 1.**
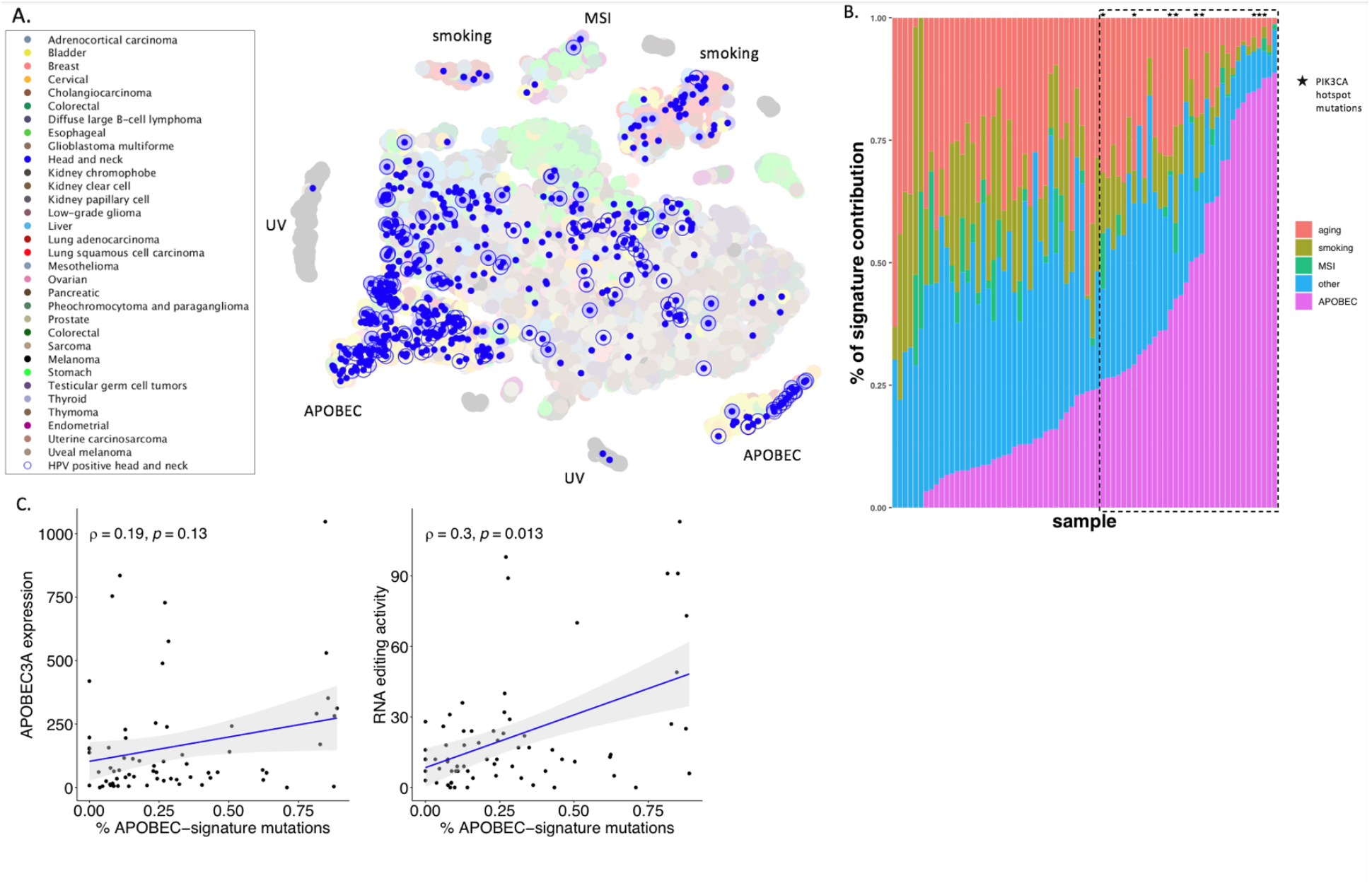
APOBEC is the dominant mutational signature in HPV+OPSCC. (A) Somatic mutation lists were combined with all cancers in TCGA, and mutational signatures were deciphered using NMF. Mutational patterns were then projected into two-dimensional space with t-SNE. Head and neck squamous cell carcinoma tumors were colored in blue dots and HPV+OPSCCs were highlighted with blue circles. Key for remaining tumor types can be found in Supplemental Figure 1A. (B) Mutational signatures ordered by APOBEC contribution (magenta) in HPV+ OPSCC samples. PIK3CA hotspot mutations, E542K and E545K, were exclusively detected within APOBEC-enriched (>25%; dashed line box) tumors. (C) Weak correlations between APOBEC mutation burden and APOBEC3A (A3A) expression (left) and A3A-associated RNA editing activity (right).

### APOBEC mutations in the viral genomes

APOBEC mutations were annotated in the HPV16 genomes after first assigning each viral genome to a sublineage. A1 (n= 45) was the predominant sublineage, followed by A2 (n=19), D3 (n=4), A4 (n=3) and C1 (n=1). There was an average (range) of 5 (0-29) mutations per viral genome. Mean (range) APOBEC mutations in the viral genomes was 1 (0-5) with an APOBEC to non-APOBEC mutation ratio mean of 1:5. Variants were most heavily clustered in the E2 C-terminal DNA-binding domain (Q349E, S364C, S348C, E344Q), including in a position previously described to impact transcription and replication of the viral genome (D338N) (Figure 2A, B)^29^. Similar to cervical HPV16 genomes^23^, there was no clear strand bias for mutations, as previously described in somatic genomes^30^. APOBEC mutations were found predominantly at very low and very high VAFs (Figure 2C). Viral APOBEC mutations, compared to non-APOBEC mutations, were more likely to be low-VAF mutations (occurring newly within the host) while non-APOBEC mutations were more likely to be high-VAF (existing prior to infection of the current host), suggesting that APOBEC mutagenesis is actively occurring in viral genomes in the current host. Viral APOBEC mutations had a higher nonsynonymous/synonymous ratio while non-APOBEC mutations showed depletion of non-synonymous variants at high VAFs compared to expectation, suggesting the influence of purifying selection (Figure 2E).

**Figure 2.**
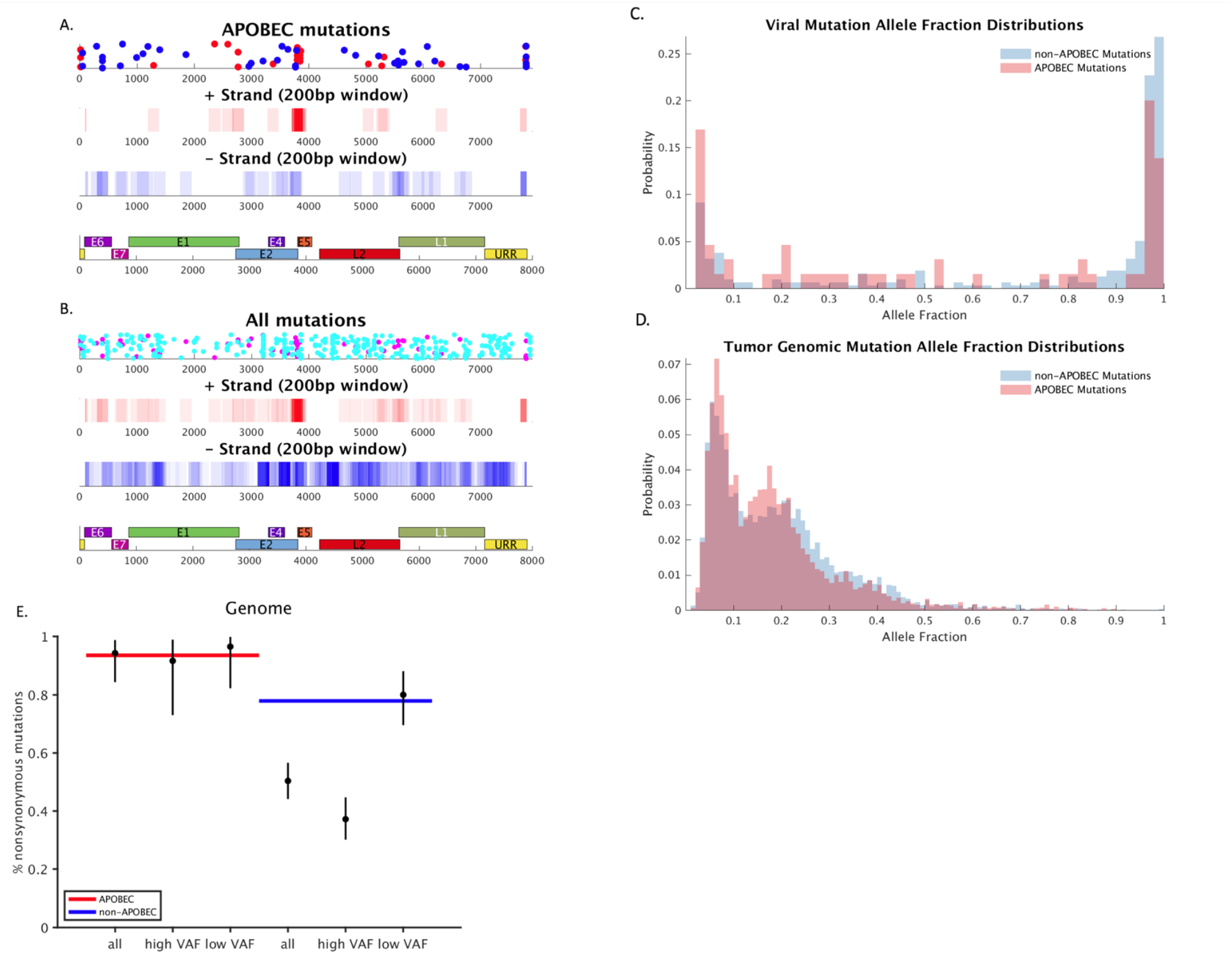
Mutation distribution in the viral genomes. APOBEC (A) and all viral mutations (B) distributed across the genome with positive strand mutations in red and negative strand in blue, highlighting enrichment of variants in E2 (panel four). Second and third panel represent mutation density on positive and negative strand with 200 base pair window. Viral (C) and tumor (D) APOBEC and non-APOBEC mutations by variant allele fraction demonstrating predominance of very low and very high VAF mutations in the viral genome. (E) Expected (blue and red lines) vs actual (black dots) nonsynonymous/synonymous ratio for APOBEC (red) and non-APOBEC (blue) mutations demonstrating higher non-synonymous/synonymous ratios for APOBEC mutations and depletion of non-synonymous variants at high VAFs compared to expected in non-APOBEC mutations, suggesting purifying selection.

### Paired APOBEC analysis

To assess the relationship between APOBEC mutations in the viral and host genomes, samples were divided into APOBEC-low and -high mutational burden groupings based on host somatic mutations (Figure 3A). APOBEC-high samples had higher viral APOBEC mutation rates (*p*=0.028), suggesting that APOBEC mutagenesis in host and viral genomes are linked (Figure 3B). We further examined additional measures of APOBEC activity in both tumor and virus and found similar trends for APOBEC RNA editing at the DDOST hairpin hotspot (*p*=0.008), A3A mRNA expression (*p*=0.14), and IFN-γ scores (*p*=0.053), which we and others have shown are tied to APOBEC activity (Figure 3C, D, E).

**Figure 3.**
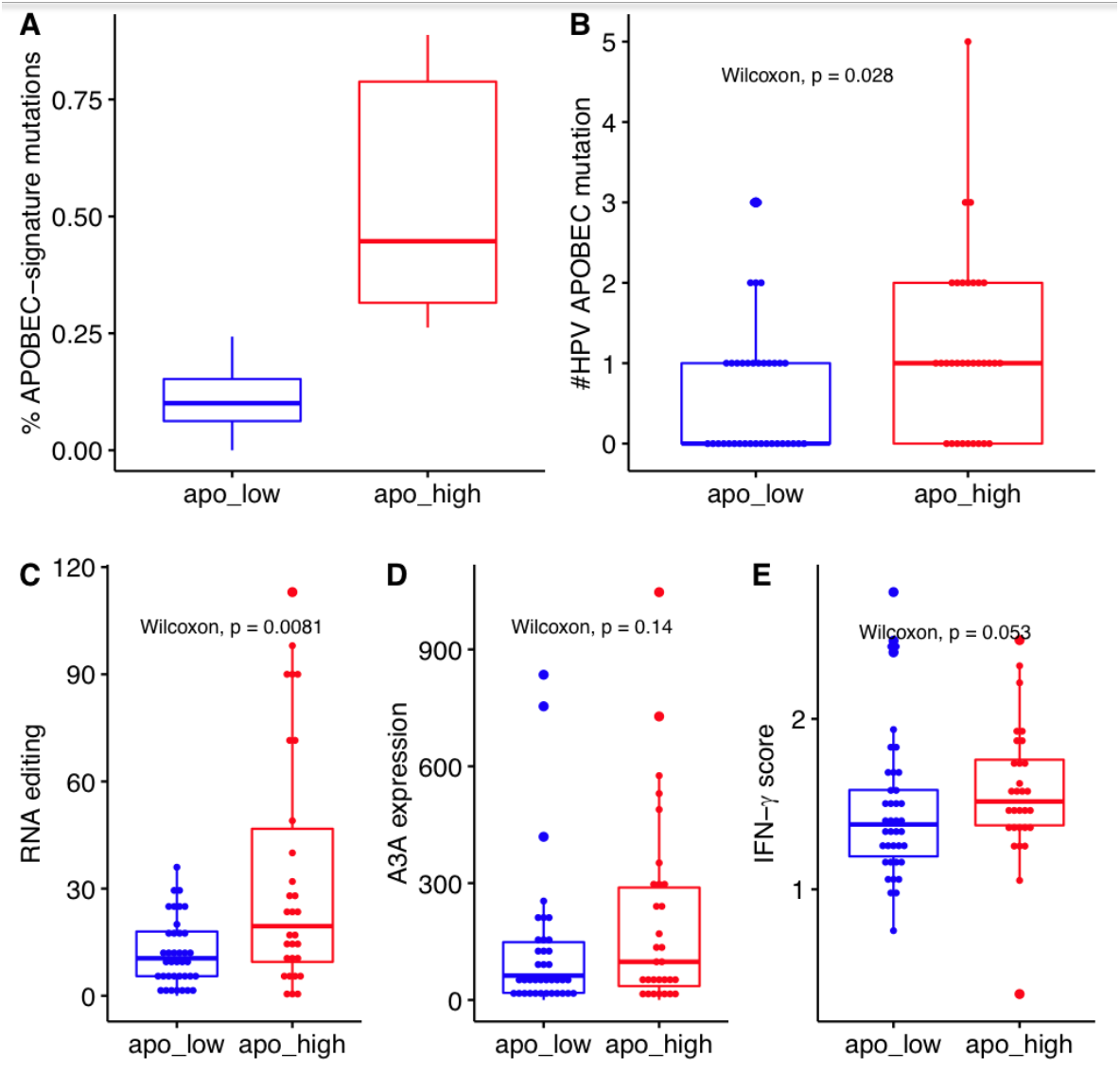
APOBEC mutations are concordant between tumor and viral genomes. APOBEC (A) mutations in tumor genomes, (B) mutation in the viral genome, (C) RNA editing and (D) 3A gene expression higher, grouped by high and low APOBEC mutational burden in the somatic genome, demonstrating linkage between APOBEC activity in virus and host. (E) APOBEC-enriched subjects also had higher IFN-γ scores, suggesting immune upregulation, possibly in response to viral infection.

## Discussion

We have conducted the first study to-date evaluating APOBEC-induced mutations in HPV+OPSCC and HPV16 genomes, demonstrating that host and viral mutations are correlated, providing a missing link connecting APOBEC mutagenesis in host and virus and supporting a common mechanism driving APOBEC dysregulation.

The development of computational approaches to determine mutational processes active in cancer genomes has led to fundamental insights into cancer development^19,31^. One such finding is that mutations occurring in the TCW context, associated with APOBEC cytidine deaminase activity, are some of the most common mutations across all cancer types^21^. While previous *in vitro* studies recognized the mutagenic potential of APOBECs in cancer genomes, the full scope of APOBEC’s mutagenic prowess was not realized until computational approaches were applied to large NGS datasets, such as TCGA^10,19^. Of particular interest was the identification that HPV-mediated cancers, of which HPV+OPSCC is the most prevalent, have the highest APOBEC mutational burdens, on average^1,19-21,31^. This finding reinforced existing *in vitro* data showing that APOBEC was active in genomes from HPV-mediated cancers and pre-cancers, and in the HPV genome itself^10^. Taken together, mounting evidence supports a role for APOBEC mutagenesis in HPV-mediated cancer genomes, regulated by mechanisms related to HPV infection.

Our group has previously characterized APOBEC-induced mutations in over 5,000 cervical benign, pre-cancer and cancer biopsies^23^, expanding on previous work and providing further support to APOBEC’s ability to mutate both cancer and viral genomes. While considerable progress has been made in quantifying the burden of APOBEC mutations in HPV-mediated cancer genomes, and cervical HPV genomes, many fundamental questions remain, including what underlying process(es) are driving APOBEC dysregulation in HPV-mediated cancers, and if APOBEC mutations in cancer and viral genomes are directly linked, or arise through different processes. Here, we utilized paired sequencing of HPV16 and host genomes to determine if APOBEC-induced mutations are present in HPV16 genomes from HPV+OPSCCs and if APOBEC activity is concordant in host and viral genomes.

Applying high-depth viral WGS, we identified APOBEC-signature mutations in OPSCC HPV16 genomes, which existed at both very low (new within host mutations) and very high (old mutations carried from prior host infections) VAFs. Viral APOBEC mutations were most prominent at low VAFs, which is of interest as HPV genomes have long been considered stable during persistent infection, yet here, and in our previous study of cervical HPV16^23^, the data suggest that APOBEC editing of viral genomes is actively occurring during infection. This is despite the fact that HPVs have evolved to have fewer APOBEC target sequence sites, likely in an effort to avoid APOBEC restriction and thus viral damage and clearence^32^. This evolutionary pattern also results in the enrichment of potential non-synonymous APOBEC mutations compared to non-APOBEC mutations. In accordance with this, we observed a high nonsynonymous-to-synonymous ratio for viral APOBEC mutations overall. The most dense cluster of APOBEC mutations occurred in E2 and includes D338N, an HPV16 SNP that has previously been reported to reduce p53 binding^29^. Interestingly, D338N is also prevalent in cervical cancer samples from our previous study. We also found depletion of nonsynonymous high-VAF mutations for non-APOBEC mutations, but did not observe this for APOBEC mutations. This could suggest that some nonsynonymous viral APOBEC-induced mutations may be beneficial to the virus and contribute to evasion of host immunity by altering viral antigens^3^. It is also possible that deleterious APOBEC mutations that result in viral damage and clearance are simply not detected as those viruses have already been cleared during initial infection stages, leaving only viral genomes with a selective growth advantage present in the cancer.

Importantly, we found that APOBEC-enriched somatic samples (greatest APOBEC mutational burden and highest level of RNA editing), had higher viral APOBEC mutation rates, providing direct evidence that APOBEC mutagenesis in host and viral genomes are linked. This suggests that in HPV-mediated cancers the underlying processes driving APOBEC dysregulation and resultant TCW context mutations, may be the same for both viral and somatic mutations. Mechanisms that drive APOBEC mutagenesis in HPV+OPSCC, however, remain unknown. While APOBEC is known to be activated by HPV oncoproteins, and through IFN signaling as part of innate immunity/viral nucleic acid sensing, APOBEC mutations are prevalent in non-virally mediated tumors, such as bladder, breast and lung cancer, as well as non-HPV mediated head and neck cancers, albeit to a lesser degree. Thus, it is possible that both viral/innate immunity and immune-independent processes, such as ssDNA targeting of the lagging strand during replication stress or after double strand breaks, drive APOBEC expression in HPV+OPSCC, and the high rates of APOBEC mutations in HPV+OPSCCs are a resultant cumulative effect. This concept is supported by work from our group and others showing multiple sources of APOBEC upregulation likely contribute to APOBEC mutagenesis in HPV+OPSCC including APOBEC germline polymorphisms and immune upregulation in response to mutation-induced neoantigens^1^.

Studies of APOBEC mutagenesis in HPV+OPSCC are limited by the lack of model systems to recapitulate early stages of HPV infection in the oropharynx and HPV+OpSCC development. Further, unlike cervical cancer, which has well defined and identifiable stages of benign infection, pre-malignancy and cancer, HPV+OPSCC has no such intermediaries. Thus murine and *in vitro* studies aimed at revealing what processes drive APOBEC mutations in HPV+OPSCC are limited by an inability to accurately recapitulate what we observe in humans. The development of accurate model systems is needed to help elucidate the role of APOBEC mutations in HPV+OpSCC, and the processes driving APOBEC dysregulation. Lastly, the relationship between ApOBEC mutational burden, tumor behavior and clinical outcomes remains to be elucidated in HPV+OPSCC.

In summary, we report the first study evaluating APOBEC mutagenesis in HPV+OPSCC somatic and viral genomes, identifying the presence of APOBEC mutations in OPSCC HPV genomes and concordance between APOBEC mutational burden within virus and host. These data provide a missing link connecting APOBEC mutagenesis in host and virus and support a common mechanism driving APOBEC dysregulation.

